# Macaque amygdala, claustrum and pulvinar support the cross-modal association of social audio-visual stimuli based on meaning

**DOI:** 10.1101/2022.09.28.509981

**Authors:** Mathilda Froesel, Maëva Gacoin, Simon Clavagnier, Marc Hauser, Quentin Goudard, Suliann Ben Hamed

**Affiliations:** Institut des Sciences Cognitives Marc Jeannerod, UMR5229 CNRS Université de Lyon, 67 Boulevard Pinel, 69675 Bron Cedex, France; Risk-Eraser, LLC, PO Box 376, West Falmouth, MA, 02574, USA

**Keywords:** Audio-visual, multisensory, pulvinar, amygdala, claustrum, fMRI, macaque, socioemotional

## Abstract

Social communication draws on several cognitive functions such as perception, emotion recognition and attention. In a previous study, we demonstrated that macaques associate audiovisual information when processing their species-specific communicative signals. Specifically, cortical activation is inhibited when there is a mismatch between vocalisations and social visual information whereas activation is enhanced in the lateral sulcus, superior temporal sulcus as well as a larger network composed of early visual and prefrontal areas when vocalisations and social visual information match. Here, we use a similar task and functional magnetic resonance imaging to assess the role of subcortical structures. We identify three subcortical regions involved in audio-visual processing of species-specific communicative signal: the amygdala, the claustrum and the pulvinar. Like the cortex, these subcortical structures are not activated when there is a mismatch between visual and acoustic information. In contrast, the amygdala and claustrum are activated by visual, auditory congruent and audio-visual stimulations. The pulvinar responds in a task-dependent manner, along a specific spatial sensory gradient. Anterior pulvinar responds to auditory stimuli, medial pulvinar is activated by auditory, audio-visual and visual stimuli and the dorsal lateral pulvinar only responds to visual stimuli in a pure visual task. The medial pulvinar and the amygdala are the only subcortical structures integrating audio-visual social stimuli. We propose that these three structures belong to a multisensory network that modulates the perception of visual socioemotional information and vocalizations as a function of the relevance of the stimuli in the social context.

**Significance Statement:** Understanding and correctly associating socioemotional information across sensory modalities, such that happy faces predict laughter and escape scenes screams, is essential when living in complex social groups. Using functional magnetic imaging in the awake macaque, we identify three subcortical structures – amygdala, claustrum and pulvinar - that only respond to auditory information that matches the ongoing visual socioemotional context, such as hearing positively valenced coo calls and seeing positively valenced grooming monkeys. We additionally describe task-dependent activations in the pulvinar, organizing along a specific spatial sensory gradient, supporting its role as a network regulator.

## Introduction

In a wide variety of species, social communication often involves sending and receiving systems that must integrate information across different modalities, recruiting multiple processes such as sensory perception, emotion processing and attention. In a previous study, we demonstrate thanks to cardiac recordings (Froesel et al., 2020) and functional magnetic resonance imaging (fMRI) that macaques associate communicatively salient audio-visual information based on the social context set by visual information. This process is mediated by a network of face and voice patches in the superior temporal sulcus and lateral sulcus (Froesel et al., 2022). Here, using the same task and data, we identify the relevant subcortical structures in cross-modal association.

Given that the task involves functionally meaningful communicative signals that are emotionally salient, we predicted strong amygdala activation. This subcortical nucleus contains face-selective neurons that are globally activated both by face identity and facial expressions (Nakamura et al., 1992; Fitzgerald et al., 2006; Pessoa et al., 2006; Sergerie et al., 2008; Livneh et al., 2012; Todorov, 2012). Specifically, affiliative facial expressions induce a decrease of firing rates in the amygdala, while threatening faces induce an increase in firing rates (Gothard et al., 2007). Overall, beyond face processing, the amygdala is part of a social perception network and is proposed to play a central role in emotional encoding during complex social interactions (Leonard et al., 1985; Barraclough and Perrett, 2011), including during cross-modal sensory emotional processing (Dolan et al., 2001; Kuraoka and Nakamura, 2007). In addition to the amygdala, the pulvinar, the largest nucleus of the thalamus, is also involved in face processing, emotion regulation and multisensory integration (Moeller et al., 2008; Pessoa, 2010a; Froesel et al., 2021). Last, the claustrum is also activated following face patches stimulation and is proposed to play a role in face perception (Moeller et al., 2008). We thus hypothesize that all of these sub-cortical structures are activated by visual and auditory social stimuli and may play a part in multisensory integration. Social perception also involves integrating contextual, behavioural, and emotional information (Ghazanfar and Santos, 2004; Freiwald, 2020). We thus predict that the audio-visual association of social information is impacted by the functional context and mediated by both these sub-cortical structures.

Here, we describe fMRI study in awake macaques performing a passive audio-visual task manipulating the congruency of cross-modal social information across six different emotional contexts, together with a passive visual task manipulating monkey facial expressions. We show, in parallel to our previous report of cortical activation, that sub-cortical activations are determined by whether monkey vocalisations match or mismatch the faces or actions associated with them. As predicted, the subcortical structures identified included the amygdala, claustrum and pulvinar. As we reported in cortical activations, these regions are modulated by the task context. Auditory activations are significant for auditory stimuli that are congruent with the visual context, but not for incongruent stimuli, indicating that inhibitory or facilitatory feedback on the context filtering auditory stimuli extends to subcortical areas. We further report three novel observations. First, the amygdala and the claustrum respond to all of visual, auditory and audio-visual stimuli, demonstrating their involvement in audio-visual integrative processes. However, only the amygdala demonstrates multisensory integration. Second, the pulvinar is activated along a sensory gradient that is determined by stimulation modality (visual, auditory or audio-visual) or context. Anterior pulvinar is only activated by auditory stimulations. Medial pulvinar is activated by visual, auditory and audio-visual congruent stimuli. The dorsal lateral pulvinar is only activated by visual stimuli in the pure visual task (For review, see Froesel et al., 2021). Third, the functional activations observed in the medial pulvinar are similar to those observed in the amygdala and claustrum, thus suggesting a coordinated role of these three sub-cortical structures in multisensory social processing. Medial pulvinar is also the only pulvinar subregion that expresses multisensory integration. Overall, our observations shed a new light on how functionally meaningful, multimodal social signal are processed by subcortical structures.

## Material and methods

### Subjects and surgical procedures

Two male rhesus monkeys (*Macaca mulatta*) participated in the study (T, 15 years, 10kg and S, 12 years, 11kg). The animals were implanted with a Peek MRI-compatible headset covered by dental acrylic. The anaesthesia for the surgery was induced by Zoletil (Tiletamine-Zolazepam, Virbac, 5 mg/kg) and maintained by isoflurane (Belamont, 1–2%). Post-surgery analgesia was ensured thanks to Temgesic (buprenorphine, 0.3 mg/ml, 0.01 mg/kg). During recovery, proper analgesic and antibiotic coverage was provided. The surgical procedures conformed to European and National Institutes of Health Guidelines for the Care and Use of Laboratory Animals. The project was authorized by the French Ministry for Higher Education and Research (project no. 2016120910476056 and 1588-2015090114042892) in accordance with the French transposition texts of Directive 2010/63/UE. This authorization was based on ethical evaluation by the French Committee on the Ethics of Experiments in Animals (C2EA) CELYNE registered at the national level as C2EA number 42.

### Experimental setup

During the scanning sessions, monkeys sat in a sphinx position in a plastic monkey chair (Vanduffel et al., 2001) facing a translucent screen placed 60 cm from the eyes. Visual stimuli were retro-projected onto this translucent screen. Their head was restrained and the auditory stimuli were displayed by Sensimetrics MRI-compatible S14 insert earphones. The monkey chair was secured in the MRI with safety rubber stoppers to prevent any movement. Eye position (X, Y, right eye) was recorded thanks to a pupil-corneal reflection video-tracking system (EyeLink at 1000 Hz, SR-Research) interfaced with a program for stimulus delivery and experimental control (EventIDE®). Monkeys were rewarded for maintaining fixation into a 2×2° tolerance window around the fixation point.

### General audio-visual run design

On each run, monkeys were required to fixate a central cross on the screen (Figure 1A). Runs followed a block design. Each run started with 10 s of fixation in the absence of sensory stimulation followed by three repetitions of a pseudo-randomized sequence containing six possible 16 s blocks: fixation (Fx), visual (Vi), auditory congruent (AC), auditory incongruent (AI), congruent audio-visual (VAC) and incongruent audio-visual (VAI). Each block (except the fixation block) consisted of alternating 500 ms stimuli (except for lipsmacks, 1s dynamic stimuli succession) of the same semantic category (see Stimuli section below), in the visual, auditory or audio-visual modalities. Each block ended with 10 s of fixation in the absence of sensory stimulation. Note that within any one run the visual stimulations were prevalent and always reflecting the same emotional content, thus setting the emotional context of the run. The initial blocks always contained a visual stimulation (V, VAC or VAI) such that pure auditory blocks could be defined as congruent or incongruent relative to the visual context set by previous blocks.

**Figure 1.**
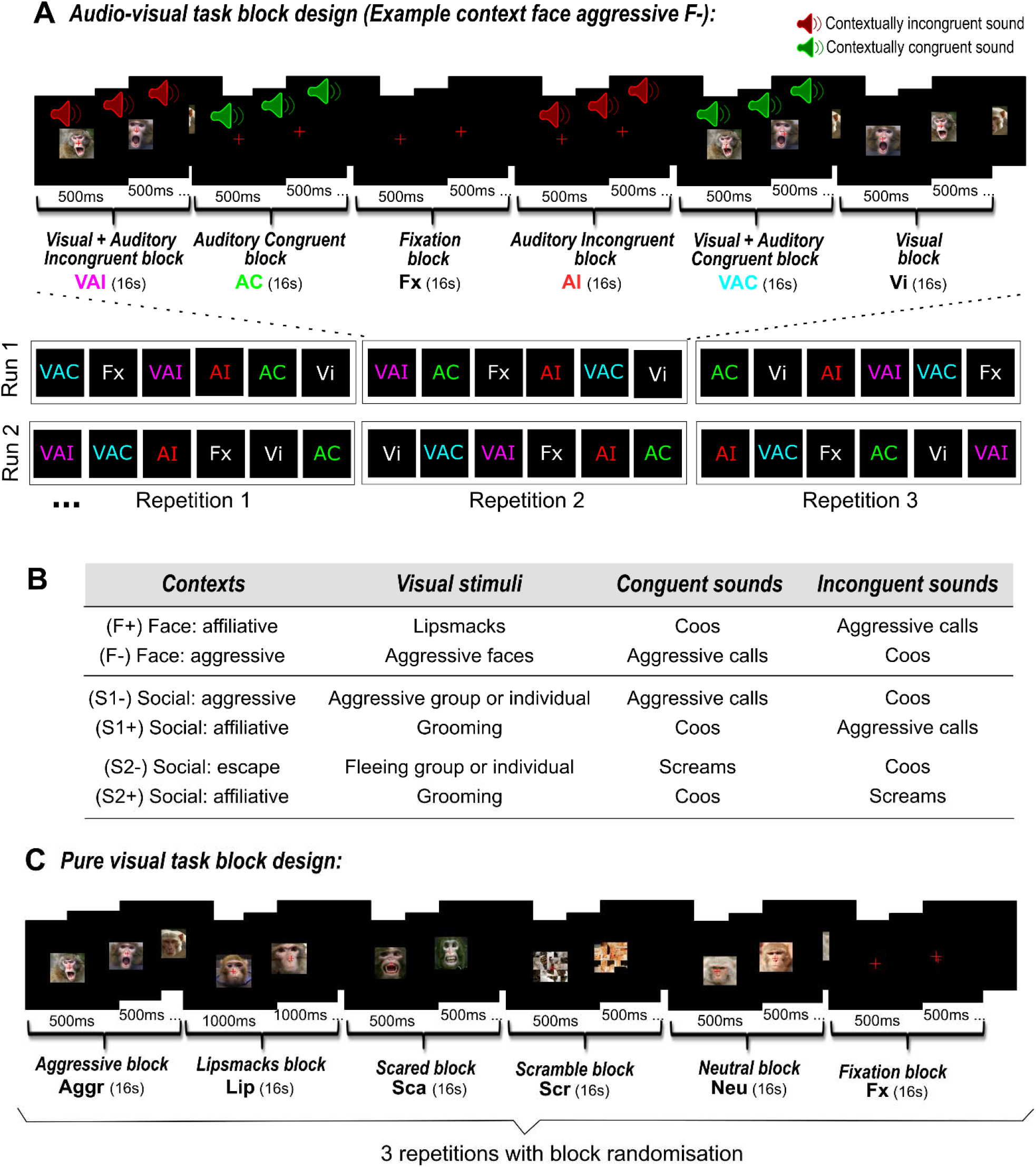
Experimental designs. Each sensory stimulation block contained a rapid succession of 500 ms stimuli (with the exception of the lipsmack for the pure visual task). Each run started and ended with 10□s of fixation regardless of the task type. **A**) Experimental design of the audio-visual task. Example of an aggressive face (F-) context. One run represents one context set up by the visual stimuli and contains three randomized repetitions of six different blocks of 16□s. The six blocks displayed were either visual stimuli only (Vi), auditory congruent stimuli only (AC), auditory incongruent stimuli only (AI), audio-visual congruent stimuli (VAC) or audio-visual incongruent stimuli (VAI), or fixation with no sensory stimulation (Fx). Blocks were pseudo randomized in order that each block was, on average, preceded by the same number of blocks from the other conditions and that each run started with a block of a visual information (V, VAC or VAI). **B**) Description of the contexts. Six contexts were displayed. Each context combined visual stimuli of identical social content with either semantically congruent or incongruent monkey vocalisations. Pairs of contexts shared the same auditory stimuli, but opposite social visual content (F+ vs. F−; S1+ vs. S1−; S2+ vs. S2−). Each run corresponded to one of the semantic contexts described above. **C**) Experimental design of the pure visual task. In one run six different blocks were displayed three times randomly. The six possible 16 s blocks were: fixation (Fx), lipsmack (Lip), scared monkey faces (Sca), aggressive monkey faces (Aggr), neutral monkey faces (Neu) and scrambled monkey faces (Scr). Visual stimuli were extracted from videos collected by the Ben Hamed lab, as well as by Marc Hauser on Cayo Santiago, Puerto Rico.

### Face and social task design

Six audio-visual contexts were presented to both monkeys, organized in runs as described above (Figure 1B). Each run combined visual stimuli of identical social content with either semantically congruent or incongruent monkey vocalisations (Figure 1B). The face affiliative context (F+) combined lipsmacks with coos and aggressive calls. The face aggressive context (F−) combined aggressive faces with coos and aggressive calls. The first social affiliative context (S1+) combined grooming scenes with coos and aggressive calls. The second social affiliative context (S2+) combined grooming scenes with coos and screams. The social aggressive context (S1−) combined aggressive group or individual scenes with coos and aggressive calls. The social escape context (S2−) combined fleeing groups or individual scenes with coos and screams. Importantly, pairs of contexts (F+ &. F−; S1+ & S1−; S2+ & S2−) shared the same auditory conditions, but opposite social visual content.

### Pure Visual runs design

The design of the visual runs was similar to that of the audio-visual run design, organized in blocks, except for the fact that all blocks were pure visual blocks and varied as a function of facial emotions. The six possible 16 s blocks were: fixation (Fx), lipsmack (Lip), scared monkey faces (Sca), aggressive monkey faces (Aggr), neutral monkey faces (Neu) and scrambled monkey faces (Scr). As for the audio-visual runs each block consisted in an alternation of 500 ms stimuli (except for lip smacks, 1s dynamic stimuli succession) of the same emotional category (see Figure 1C).

### Stimuli

Vocalisations were recorded by Marc Hauser from semi-free-ranging rhesus monkeys during naturally occurring situations. Detailed acoustic and functional analyses of this repertoire has been published elsewhere (e.g., Gouzoules et al., 1984; Hauser & Marler, 1993). Field recordings were then processed, restricting selection of experimental stimuli to calls that were recorded from known individuals, in clearly identified situations, and that were free of competing noise from the environment. Exemplars from this stimulus set have already been used in several imaging studies (Belin et al., 2007; Cohen et al., 2007; Romanski, 2012; Romanski et al., 2005; Russ et al., 2008). As in our previous study (Froesel et al., 2022), all stimuli were normalized in luminance and colour but the frequency ranges varied between the different types of stimuli as shown in the supplementary Figure 4. For each of the three vocalisation categories, we used 10 unique exemplars coming from matched male and female individuals for each category in order to control for possible gender, social hierarchy or individual effects. Coos are affiliative vocalisations, aggressive calls are used as a precursor of a physical attack and screams are produced by subordinate being chased or attacked by a dominant. Facial expression (lipsmacks and aggressive facial expression) and social scene (group grooming, aggressive individual alone or in group / escaping individual or group) stimuli were extracted from videos collected by the Ben Hamed lab, as well as by Marc Hauser on Cayo Santiago, Puerto Rico. Images were normalized for average intensity and size. All stimuli were 4° x 4° in size. We decided to keep them in colour to get closer to natural stimuli even if it produced greater luminosity disparity between the different stimuli preventing us to use pupil diameter as a physiological marker. Only unambiguous facial expressions and social scenes were retained. A 10% blur was applied to all images, in the hope of triggering multisensory integration processes (Stein and Meredith, 1993)(but see result section). For each visual category, 10 stimuli were used. The scrambling of the images was performed by EventIDE (https://www.okazolab.com/okazolab.com/) and applied to each visual stimuli displayed from the task.

### Scanning Procedures

The in-vivo MRI scans were performed on a 3T Magnetom Prisma system (Siemens Healthineers, Erlangen, Germany). For the anatomical MRI acquisitions, monkeys were first anesthetized with an intramuscular injection of ketamine (10 mg\kg). Then, the subjects were intubated and maintained under 1-2% of isoflurane. During the scan, animals were placed in a sphinx position in a Kopf MRI-compatible stereotaxic frame (Kopf Instruments, Tujunga, CA). Two L11 coils were placed on each side of the skull and a L7 coil was placed on the top of it. T1-weighted anatomical images were acquired for each subject using a magnetization-prepared rapid gradient-echo (MPRAGE) pulse sequence. Spatial resolution was set to 0.5 mm, with TR= 3000 ms, TE=3.62 ms, Inversion Time (TI)=1100 ms, flip angle=8°, bandwidth=250 Hz/pixel, 144 slices. T2-weighted anatomical images were acquired per monkey, using a Sampling Perfection with Application optimized Contrasts using different flip angle Evolution (SPACE) pulse sequence. Spatial resolution was set to 0.5 mm, with TR= 3000 ms, TE= 366.0 ms, flip angle=120°, bandwidth=710 Hz/pixel, 144 slices. fMRI acquisitions were as follows. Before each scanning session, a contrast agent, composed of monocrystalline iron oxide nanoparticles, Molday ION™, was injected into the animal’s saphenous vein (9-11 mg/kg) to increase the signal to noise ratio (Vanduffel et al., 2001; Leite et al., 2002). We acquired gradient-echoechoplanar images covering the whole brain (TR=2000 ms; TE=18 ms; 37 sagittal slices; resolution: 1.25×1.25×1.38 mm anisotropic voxels) using an eight-channel phased-array receive coil; and a loop radial transmit-only surface coil (MRI Coil Laboratory, Laboratory for Neuro- and Psychophysiology, Katholieke Universiteit Leuven, Leuven, Belgium, see Kolster et al., 2014). The coils were placed so as to maximise the signal on the temporal lobe.

### Data description

In total, for the audio-visual runs (155 pulses), 76 runs were collected in 12 sessions for monkey T and 65 runs in 9 sessions for monkey S. For the pure visual runs (155pulses), 13 runs were collected in 8 sessions for monkey T and 12 runs in 5 sessions for monkey S. Based on the monkey’s fixation quality during each run (85% within the eye fixation tolerance window), we selected 60 runs from monkey T and 59 runs for monkey S in total, i.e. 10 runs per task, except for one task of monkey S for audio-visual runs and 11 pure visual runs for S and 13 for T.

### Data analysis

Data were pre-processed and analysed using AFNI (Cox, 1996), FSL (Jenkinson et al., 2012; Smith et al., 2013), SPM software (version SPM12, Wellcome Department of Cognitive Neurology, London, UK, https://www.fil.ion.ucl.ac.uk/spm/software/), JIP analysis toolkit (http://www.nitrc.org/projects/jip) and Workbench (https://www.humanconnectome.org/software/get-connectome-workbench). The T1-weighted and T2-weighted anatomical images were processed according to the HCP pipeline (Glasser et al., 2013; Autio et al., 2020) and were normalized into the MY19 Atlas (Donahue et al., 2016). Functional volumes were corrected for head motion, slice timed referred on the middle image of the run and skull-stripped. They were then linearly realigned on the T2-weighted anatomical image with flirt from FSL, the image distortions were corrected using nonlinear warping with JIP. A spatial smoothing was applied with a 3-mm FWHM Gaussian Kernel.

Fixed effect individual analyses were performed for each monkey, with a level of significance set at p<0.05 corrected for multiple comparisons (FWE, t-scores 4.6) and p<0.001 (uncorrected level, t-scores 3.09). Head motion and eye movements were included as covariate of no interest. Because of the contrast agent injection, a specific MION hemodynamic response function (HRF) (Vanduffel et al., 2001) was used instead of the BOLD HRF provided by SPM. The main effects were computed over both monkeys. In most analyses, face blocked conditions and social blocked conditions were independently pooled.

ROI analyses were performed as follows. The anterior pulvinar, claustrum ROIs were determined from the auditory congruent contrast (AC vs Fx) and the medial pulvinar and the amygdala ROIs from the audio-visual contrast (VAC vs FX) of face context (Figure 2). PLvl and Pldm ROIs were extracted from the pure visual runs (all conditions vs fixation contrast; figure 4). ROIs were defined as 1 mm diameter spheres centred around the local peaks of activation. To note, there was no overlap between the different clusters. For each ROI, the activity profiles were extracted with the Marsbar SPM toolbox (marsbar.sourceforge.net) and the mean percent of signal change (+/- standard error of the mean across runs) was calculated for each condition relative to the fixation baseline. As the face context includes both aggressive and lipsmack expressions, we decided for pure visual runs to combine these two expressions conditions and focus on this combination for the analysis to be comparable to the PSC results from the audiovisual runs. %SC were compared using Friedman non-parametric tests and Wilcoxon nonparametric paired tests.

**Figure 2:**
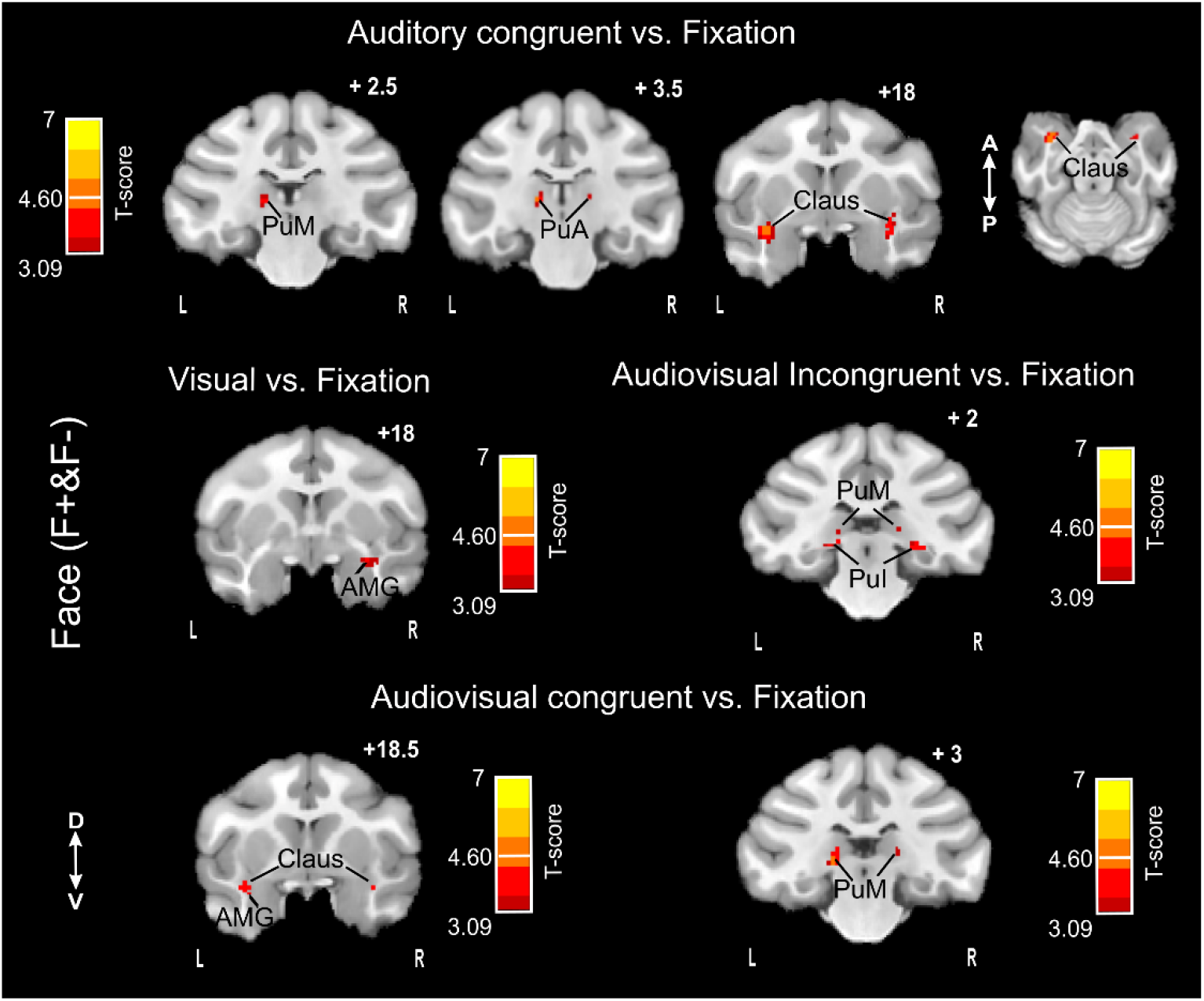
Whole-brain FACE blocked conditions (F+ & F-) activations: main contrasts. Whole-brain activation maps of the F+ (face affiliative) and F- (face aggressive) runs, cumulated over both monkeys, for the visual, auditory congruent, audio-visual congruent and audio-visual incongruent contrasts. Darker shades of red indicate level of significance at p<0.001 uncorrected, t-score 3.09. Lighter shades of yellow indicate level of significance at p<0.05 FWE, t-score 4.6.

## Results

In Froesel et al., 2022, we characterized cortical activations underlying the association of social visual and auditory stimuli based on their semantic content or meaning in the broad sense. Cortical regions were active when visual and auditory information were congruent, but inactive when this information was incongruent. Here, we identify subcortical structures -the amygdala, the claustrum and the pulvinar-that also respond to the congruence of the auditory and visual stimuli. In addition, we show that among subcortical regions, only the amygdala and the medial pulvinar implement multisensory integration.

### Sub-cortical activations: amygdala, ventral putamen and pulvinar

A small group of sub-regions are significantly activated during one or several of the unimodal or bimodal conditions presented to the monkeys in the different types of runs. Indeed, in a general contrast analysis, the claustrum (Claus) and the anterior (PuA) and medial (PuM) pulvinar nuclei are activated by the Auditory congruent vs. Fixation contrast (Figure 2, top raw). The amygdala (AMG) is activated by the Visual vs. Fixation contrast (Figure 2, middle left panel). The inferior (PuI) and medial (PuM) pulvinar nuclei are activated by the Audio-visual incongruent vs. Fixation contrast (Figure 2, middle right panel). The amygdala (AMG), claustrum (Claus) and medial (PuM) pulvinar nucleus are activated by the Audio-visual congruent vs. Fixation contrast (Figure 2, bottom raw). All of the activations are bilateral in at least one contrast except for the amygdala. Supplemental figure S1 represents the SNR map for the coronal section at the level of the pulvinar.

We defined functional ROIs based on these sub-cortical activations and we computed the percent of signal change relative to fixation (%SC) for all conditions, cumulated over identical block conditions of either the face contexts (Figure 3, left), or the social scenes contexts (Figure 3, right). For all ROIs, and both face and social contexts, there was no significant activation relative to the fixation baseline in response to the auditory incongruent blocks. All other blocks led to consistent significant activations, except for inferior and anterior pulvinar, in the visual blocks of both face and social scene contexts, and the ventral putamen, in the visual blocks of social scene contexts.

**Figure 3:**
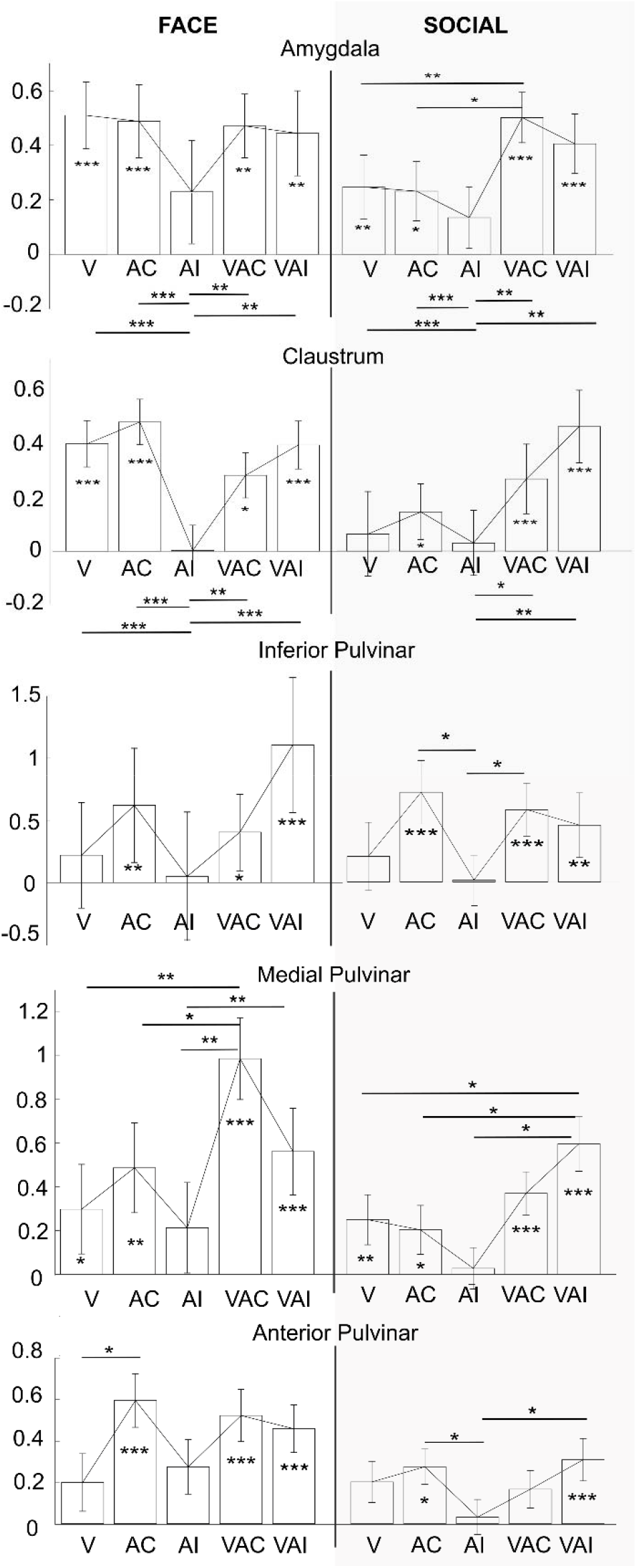
Percentage of signal change (%SC) for FACE tasks (F+ & F−) and for Social tasks (S1+, S1-, S2+ & S2-) across sub cortical ROIs, amygdala, claustrum, inferior pulvinar, medial pulvinar and anterior pulvinar of both hemispheres, comparing the auditory, visual and audiovisual conditions. Statistical differences relative to fixation and between conditions are indicates as follows: ***, p<0.001; **, p<0.01; *, p<0.05 (Wilcoxon non-parametric test). See table 1 for quantitative effect sizes. <u>AMG</u>: *FACE*: V: Z=4.8, p<0.001; AC: Z=3.79, p<0.001; AI: Z=1.17, p=0.2; VAC: Z=2.9, p=0.003; VAI: Z=2.62, p=0.009. Friedman non-parametric test, X2_(4)_ = 20.83, p<0.001, n=80, post hoc: V-AI: Z=2.16, p<0.001, AC-AI: Z=1.75, p<0.001, VAC-AI: Z=1.74, p<0.01, VAI-AI: Z=1.4, p<0.01). *SOCIAL*: V: Z=2.9, p=0.0036; AC: Z=1.9, p=0.05; AI: Z=1.03, p=0.29; VAC: Z=5.6, p<0.001; VAI: Z=4.7, p<0.001. Friedman non-parametric test, X2_(4)_ = 32.77, p<0.001, n=158; post hoc: VAC-V: Z=2.36, p=0.01; VAC-AC: Z= 2.25, p=0.022; AI-V VAC-AI: Z= 3.09, p=0.0019; VAC-AI: Z= 2.52, p=0.01). <u>Claus</u>: *FACE*:> V: Z=3.79, p<0.001; AC: Z=3.8, p<0.001; AI: Z=0, p=0.9; VAC: Z=2.04, p=0.0412; VAI: Z=4.37, p=0.009. Friedman non-parametric test, X2(_4_) = 24.83, p<0.001, n=80, post hoc: V-AI: Z=2.4, p<0.001, AC-AI: Z=2.9, p<0.001, VAC-AI: Z=1.77, p<0.01, VAI-AI: Z=2.6, p<0.01). *SOCIAL*: V: Z=1.45, p=0.14; AC: Z=2.28, p=0.022; AI: Z=0.2, p=0.83; VAC: Z=4.57, p<0.001; VAI: Z=5.40, p<0.001. Friedman non-parametric test, X2_(4)_ = 26.82, p<0.001, n=158; post hoc: VAC-AI: Z=2.23, p=0.025; VAI-AI: Z= 2.58, p=0.009). <u>PuI</u>: *FACE*:> V: Z=0.87, p=0.38; AC: Z=3.2, p=0.0013; AI: Z=1.45, p=0.14; VAC: Z=2.3, p=0.012; VAI: Z=5.25, p<0.001. Friedman non-parametric test, X2(4) = 6.48, p=0.1, n=80). *SOCIAL*: V: Z=1.24, p=0.2; AC: Z=4.78, p<0.001; AI: Z=0, p=1; VAC: Z=3.74, p<0.001; VAI: Z=2.70, p=0.006. Friedman non-parametric test, X2_(4)_ = 19.25, p<0.001, n=158, post hoc: AC-AI: Z=2.6, p=0.009; VAC-AI: Z=1.9, p=0.049). <u>PuM</u>: *FACE*:> V: Z=2.04, p=0.04; AC: Z=2.6, p<0.01; AI: Z=1.7, p=0.08; VAC: Z=5.25, p<0.001; VAI: Z=3.79, p<0.001. Friedman nonparametric test, X2(4) = 19.26, p<0.001, n=80, post hoc: V-VAC: Z=1.6, p<0.01, AC-VAC: Z=1.2, p=0.02, VAC-AI: Z=1.9, p<0.01, VAI-AI: Z=1.37, p<0.01). *SOCIAL*: V: Z=2.9, p=0.003; AC: Z=2.01, p=0.037; AI: Z=1.03, p=0.29; VAC: Z=3.95, p<0.001; VAI: Z=3.96, p<0.001. Friedman non-parametric test, X2_(4)_ = 10.54, p=0.03, n=158; post hoc: VAI-V: Z=1.86, p=0.05; VAI-AC: Z=1.9, p=0.049, VAI-AI: Z=2, p=0.048). <u>PuA</u>: *FACE*:> V: Z=1.16, p=0.24; AC: Z=4.08, p<0.001; AI: Z=1.45, p=0.14; VAC: Z=4.37, p<0.001; VAI: Z=3.79, p<0.001. Friedman non-parametric test, X2(4) = 11.59, p=0.02, n=80, post hoc: V-AC: Z=1.7, p=0.04). *SOCIAL*: V: Z=1.45, p=0.14; AC: Z=2.28, p=0.02; AI: Z=0.2, p=0.8; VAC: Z=1.87, p=0.06; VAI: Z=4.36, p<0.001. Friedman non-parametric test, X2_(4)_ = 18, p= 0.0012, n=158, post hoc: AC-AI: Z=1.88, p=0.05 VAI-AI: Z=1.9, p=0.049).

### Amygdala and claustrum audio-visual activations

In the amygdala and the claustrum, %SC in response to the auditory incongruent blocks relative to fixation, was statistically significantly lower than the %SC in all other conditions (i.e., V, AC, VAC and VAI), in both the face and the social contexts (figure 3, two top panels, except for the claustrum, in the social scene where AI was not significantly different from V and AC). The pattern of response in these two sub-cortical structures was thus very close to that reported in Froesel et al. (2022) at the cortical level. Interestingly, the amygdala shows a higher activation following audio-visual congruent stimulation, i.e. in the bimodal stimulation, than for unimodal stimulation, i.e. visual only and auditory only contexts, in the social context. This was similar to what we previously described in the superior temporal sulcus in Froesel et al. (2022), suggesting a functional interaction between these two regions.

### Pulvinar audio-visual activations

Quite unexpectedly, in the three different pulvinar ROIs (inferior, medial and anterior), although there was no significant %SC in response to the auditory incongruent condition relative to fixation, in most cases, no statistically significant difference could be observed between the %SC in this condition and the other conditions (i.e., V, AC, VAC and VAI), in neither the face context nor the social context (figure 3, three bottom panels), except for the medial pulvinar. Indeed, in medial pulvinar, marked significant differences could be observed between the %SC in the AI condition and in the VAC and/or VAI conditions in both types of context (FACE: Friedman non-parametric test, X2_(4)_ = 19.26, p<0.001, n=80, post hoc: VAC-AI: Z=1.9, p<0.01, VAI-AI: Z=1.37, p<0.01; SOCIAL: Friedman nonparametric test, X2(4) = 10.54, p= 0.03, n=158; post hoc: VAI-AI: Z=2, p=0.048).

PuM was also the only pulvinar subregion to respond to visual stimulation presented alone (*FACE*: V: Z=2.04, p=0.04; V: Z=2.9, p=0.003). In addition, and paralleling amygdala activation, the medial pulvinar showed a higher activation in bimodal (VAC) versus unimodal conditions (V and AC), specifically to the face context (FACE: Friedman non-parametric test, X2(_4_) = 19.26, p<0.001, n=80, post hoc: V-VAC: Z=1.6, p<0.01, AC-VAC: Z=1.2, p=0.02).

This leads to the noteworthy and unexpected observation that, in this task, the inferior and anterior pulvinar preferentially respond in the auditory conditions, i.e. AC, VAC and VAI conditions rather than in the visual condition, while PuM singles out responding to all conditions and exhibiting a form of high-level multisensory integration.

### Amygdala, claustrum and pulvinar activations in a purely visual task

Because macaque pulvinar has been repeatedly observed to be involved in visual processing, we hypothesized that this specific pulvinar response pattern is a task effect and reflects a spatial segregation of pulvinar sensory responses. To test this, we investigated pulvinar activations in a pure visual task, in which only monkey faces were presented in blocks of consistent emotional categories. In order to match the visual categories used in the main audio-visual task, the following, we only used the lipsmacks and aggressive categories. Clear lateralized left visual activations could be identified in the lateral pulvinar (figure 4, dorsally: PLdm and ventrally: PLvl), at a location very distinct from the ROIs activated in the audio-visual tasks. In this same task and same contrast, the claustrum was also significantly activated (figure 5A) but not the amygdala, although a %SC analysis using a priori defined ROIs extracted from the audio-visual task show that both subcortical structures are highly activated during these pure visual runs (figure 5B; AMG: Z=4.27; p<0.001; Claus: Z= 5.8, p<0.001). Overall, this indicates that while the amygdala and the claustrum are involved in the processing of social cues in both visual and audio-visual contexts, the spatial organization of the pulvinar activations vary between the two types of sensory contexts. This is explored in more details in the next section.

### Gradient of multi and unimodal pulvinar activations

The pulvinar is involved in several sensory processes (for review, see Froesel et al., 2021). How this sensory information from multiple modalities is organized remains poorly understood. In the following, we describe a spatial gradient of sensory responses within the pulvinar, from pure auditory responses (Figure 6A, red), audio-visual responses (Figure 6A&B, blue, see supplemental figure S2 for individual monkey maps), and pure visual responses (Figure 6B, green). More specifically, pure auditory activations are located in anterior pulvinar (PuA), anterior and medial to the audio-visual activations which are located in medial pulvinar (PuM), with a slight overlap between the two activated ROIs. The pure visual activations in the visual task are located in two distinct lateral pulvinar ROIs (Figure 6B, sagittal view, green), lateral to the PuM audio-visual ROIs (Figure 6B, blue), that coincide with PLvl and PLdm (Figure 4), again with a slight overlap between the audio-visual and the visual ROIs. It is worth noting that this gradient was present in each monkey (supplemental figure S2).

**Figure 4:**
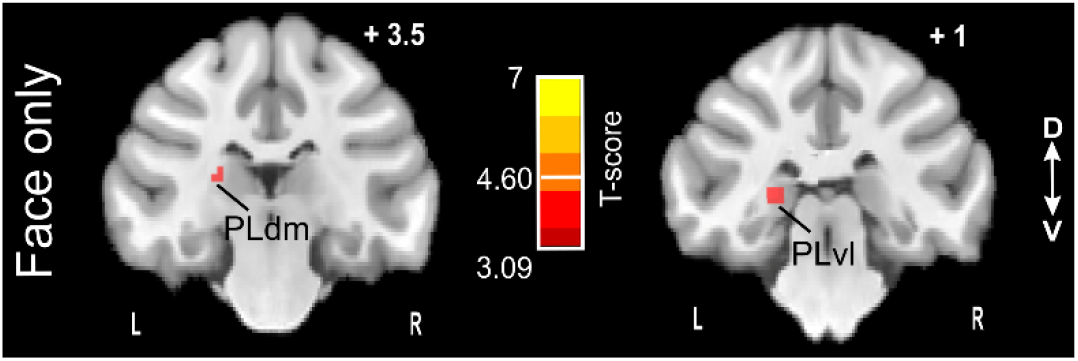
Pulvinar (and whole-brain activation) in a pure visual task: (lipsmacks + aggressive) conditions vs. fixation contrasts. Whole-brain activation maps of the pure visual task cumulated over both monkeys, for lipsmack + aggressive blocks versus fixation contrast. Darker shades of red indicate level of significance at p<0.001 uncorrected, t-score 3.09. Lighter shades of yellow indicate level of significance at p<0.05 FWE, t-score 4.6.

**Figure 5:**
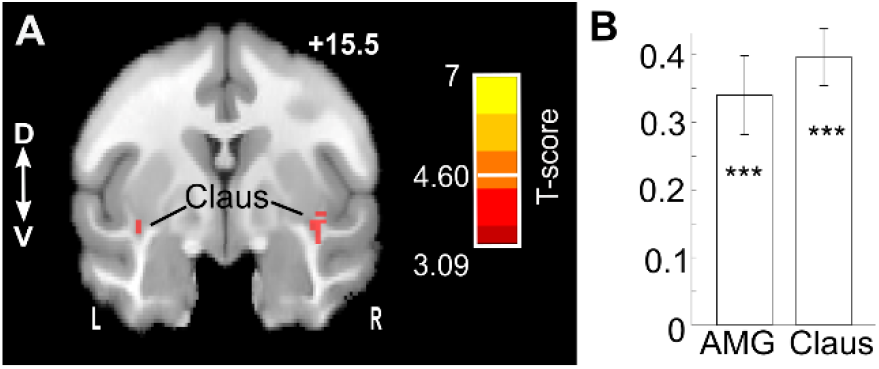
Claustrum (and whole-brain activation) in a pure visual task: (lipsmacks + aggressive) conditions vs. fixation contrasts. (A) Whole-brain activation maps of the pure visual task cumulated over both monkeys, for lipsmack + aggressive blocks versus fixation contrast. Darker shades of red indicate level of significance at p<0.001 uncorrected, t-score 3.09. Lighter shades of yellow indicate level of significance at p<0.05 FWE, t-score 4.6. (B) Percent signal change of lipsmack + aggressive blocks versus fixation contrast in the amygdala (AMG) and the claustrum (Claus) ROIs defined in the audio-visual task described in Figure 2. AMG: Z=4.27; p<0.001; Claus: Z= 5.8, p<0.001.

**Figure 6:**
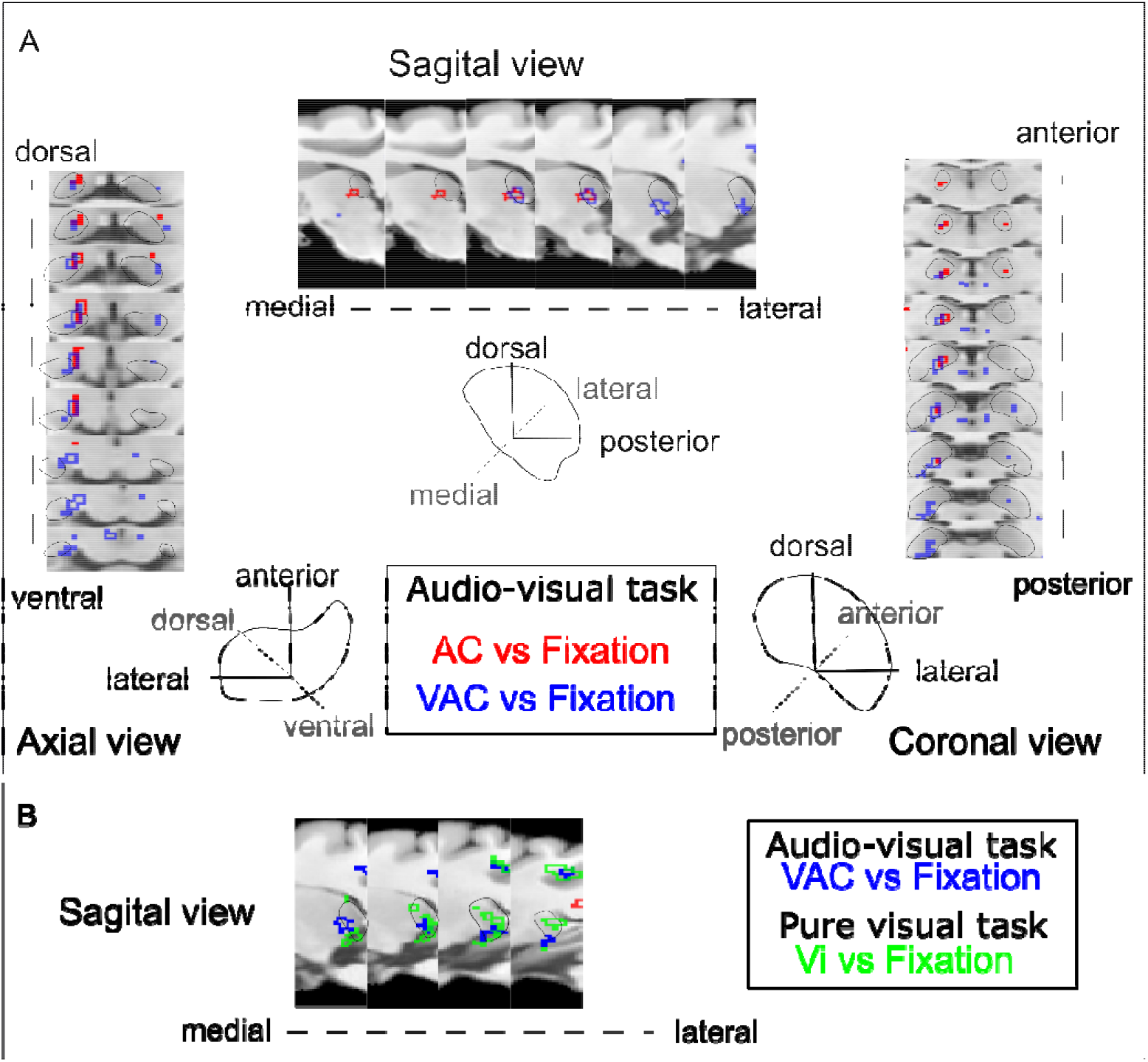
Pulvinar (and whole-brain) activations in FACE (F+ & F-) audio-visual and pure visual tasks. A) Whole-brain activation maps of the FACE context cumulated over bo**th** monkeys, for auditory congruent (red) and audio-visual congruent (blue) contrasts vs. fixatio**n**. Axial, sagittal and coronal view are shown, zooming on the pulvinar. The activation outline in r**ed** and blue correspond to activation thresholds at the level of significance p<0.001 uncorrected, t-score 3.09. B) Whole-brain activation maps of the FACE context cumulated over both monke**ys**, audio-visual congruent condition of the audio-visual task (blue) and visual condition (aggress**ive** + lipsmack faces) of the pure visual task (green).

Importantly, this very clear functional sensory gradient within the pulvinar is task specific. Indeed, the visual ROIs identified during the pure visual task are not activated during the audiovisual task using the exact same stimuli, and vice versa (Figure 7A&B). All pulvinar ROIs except the anterior pulvinar (PuA) are significantly activated relative to fixation during the pure visual task (Figure 7A; PuI: Z=3.19, p<0.001; PuM: Z=2.6, p=0.007; PuA: Z=1.86; p=0.062). In contrast, only the ventro-lateral pulvinar ROI (PLvl) retains this significant visual response in the visual only condition of the audio-visual task, although both tasks involve the same visual conditions (Figure 7B; Z=2.1, p=0.02). These results suggest that pulvinar visual responses (except for PLvl) are not fully driven by sensory input and are modulated by general task context.

**Figure 7:**
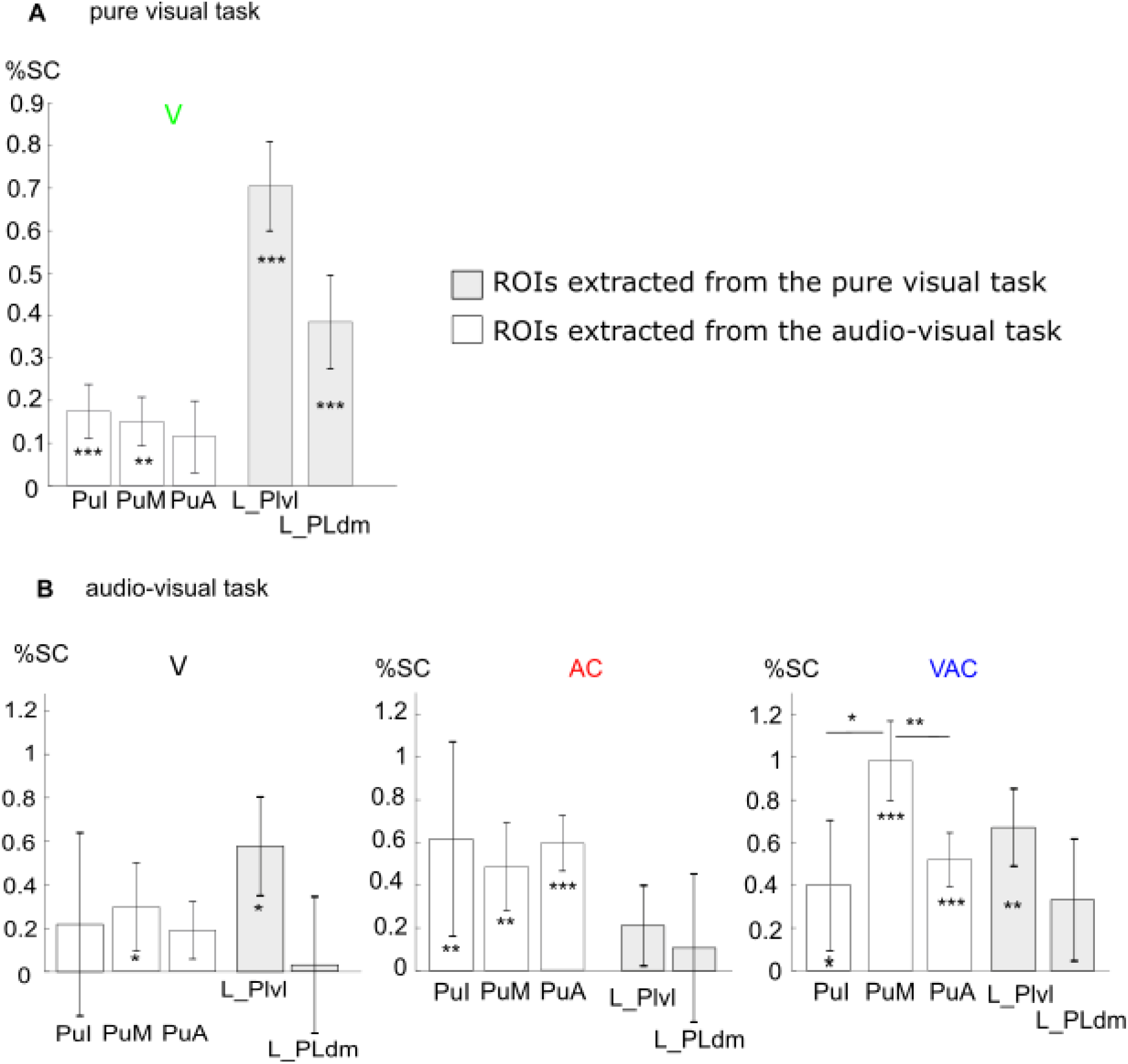
Percentage of signal change in selected pulvinar ROIs, the pure visual task (A) and the audio-visual task (B). Statistical differences relative to fixation and between conditions are indicates as follows: ***, p<0.001; **, p<0.01 (Wilcoxon non-parametric test). Selected ROIs are: Inferior pulvinar (PuI, bilateral), Medial pulvinar (PuM, bilateral), Anterior pulvinar (PuA, bilateral) and Lateral pulvinar (PLdm and PLvl, left). ROIs defined in the audio-visual task are in white. ROIs defined in the pure visual task are in gray. V: visual vs fixation; AC: auditory congruent vs fixation and VAC: audio-visual congruent vs fixation. (A) PuI: Z=3.19, p<0.001; PuM: Z=2.6, p=0.007; PuA: Z=1.86; p=0.062; PLdm: Z= 2.66, p<0.001; Plvl: Z=3.99, p<0.001. Friedman non-parametric test, X2_(4)_ = 3.94, p= 0.4, n=96. (B) <u>V</u>: PuI: Z=0.87, p=0.38; PuM: Z=2.04, p=0.04; PuA: Z=1.16, p=0.24; PLdm: Z=0.29, p=0.7; PLvl: Z=2.1, p=0.02. Friedman non-parametric test, X2_(4)_ = 7.67, p= 0.1, n=80. <u>AC</u>: PuI: Z=3.2, p=0.0013; PuM: Z=2.6, p<0.01; PuA: Z=2.6, p<0.01; PLdm: Z=0.8, p=0.38; PLvl: Z=1.16, p=0.24. Friedman non-parametric test, X2(4) = 7.87, p= 0.09, n=80. <u>VAC</u>: PuI: Z=2.3, p=0.011; PuM: Z=5.25, p<0.001; PuA: Z=4.37, p<0.001; PLdm: Z=3.2, p=0.0013; PLvl: Z=2.4, p=0.01. Friedman non-parametric test, X2_(4)_ = 9.82, p= 0.03, n=80. PuI-PuM: Z=2.1 p=0.02; PuM-PuA: Z= 2.4, p=0.01.

PLvl also retains significant %SC in the visuo-auditory condition (Figure 7B, VAC: Z=2.4, p=0.01) but not in the AC condition (Z=1.16, p=0.24), suggesting that the VAC significance is driven by the visual component of the VAC condition. Although we did not run a pure auditory task, the inferior and anterior pulvinar ROIs contrast with PLvl and appear to be driven by auditory stimulation as their %SC is significant for both the auditory (Figure 7B, AC: PuI: Z=3.2, p=0.0013; PuA: Z=2.6, p<0.01) and visuo-auditory condition (VAC: PuI: Z=2.3, p=0.011; PuA: Z=4.37, p<0.001), but not in the visual condition (V: PuI: Z=0.87, p=0.38; PuA: Z=1.16, p=0.24). Medial pulvinar ROI (PuM) stands out in that is activated by all of the visual conditions whether in the pure visual task (Figure 7A, V: Z=2.6, p=0.007) or the audio-visual task (Figure 7B, V: Z=2.04, p=0.04), the auditory condition of the audio-visual task (Figure 7B, AC: Z=2.6, p<0.01) and the audio-visual condition of the audio-visual task (Figure 7B, VAC: Z=5.25, p<0.001). Notably, the %SC in the VAC condition is significantly higher than the %SC in the V and AC conditions of the audio-visual task (Friedman non-parametric test, X2_(4)_ = 19.26, p<0.001, n=80, post hoc: V-VAC: Z=1.6, p<0.01, AC-VAC: Z=1.2, p=0.02), strongly suggesting that the medial pulvinar, PuM ROI, in contrast with the other ROIs, is integrating visual and auditory information.

## Discussion

Overall, three key subcortical structures contribute to the audio-visual association of functionally significant communicative signal: the amygdala, claustrum and pulvinar. Activation patterns in the pulvinar were not predicted based on prior research and shed new light on the functional organization of this subcortical nucleus. The medial part of the nucleus and the amygdala demonstrate multisensory integration similar to what we report in the superior temporal sulcus in a previous study using the same data (Froesel et al., 2022). It is worth noting that the amygdala, claustrum and pulvinar correspond to the three subcortical regions activated by face patch stimulations. It has been proposed that these regions correspond to a bottlenecks for the communication between face patches (Moeller et al., 2008). This interpretation is supported by a tracer studies showing that individual patches receive input from these three subcortical structures (Grimaldi et al., 2016) as well as by resting-state fMRI, alone or in association with single cell recording studies (Schwiedrzik et al., 2015; Zaldivar et al., 2022). The amygdala, claustrum and pulvinar are thus a part of a subcortical network involved in face processing. In this study, we extend their role to the processing of species-specific vocalizations and their association with socioemotional visual information. The degree of specificity is not yet established, requiring tests with other species’ vocalisations, facial expressions, and social scenes.

### Audio-visual association of social stimuli in the amygdala and the claustrum

The amygdala and claustrum are activated by all visual, auditory congruent and audio-visual (congruent and incongruent) stimuli where auditory congruence or incongruence is defined by the visual context set in each presentation. Both these subcortical structures thus follow a response pattern that is similar to that observed at the cortical level (Froesel et al., 2022). In addition, they both are also activated by emotional facial expressions during a pure visual task. This pattern suggests that both structures contribute to context-based social and emotional processing in coordination with the face and voice patches described in Froesel et al. (2022).

The contribution of the amygdala to the processing of social stimuli, whether visual (Nakamura et al., 1992; Pessoa et al., 2006; Sergerie et al., 2008; Todorov, 2012) or auditory (Gadziola et al., 2012; Domínguez-Borràs et al., 2019; Morrow et al., 2019; Gothard, 2020) has been extensively studied by others. Here, we further show that this processing is context dependent, such that a particular stimulus either activates or inactivates the amygdala depending upon the context. Additionally, we show that like the superior temporal sulcus (Froesel et al., 2022) the amygdala also plays an important role in multisensory integration (see also (Ross et al., 2022)). These activations are located in the dorsal and lateral part of the nucleus, close to the claustrum.

The claustrum is, due to its connectivity with inputs from limbic areas and many other cortical and subcortical areas, well positioned as a hub associating sensory and limbic information in order to influence attention via its output to the frontal cortex (see for review Smith et al., 2020). It has recently been hypothesized that in addition to its involvement in the coordination of slow wave sleep, it could serve as a limbic-sensory-motor interface. The claustrum is proposed to integrate limbic and sensory information to guide and sustain attention towards behaviourally relevant and salient stimuli. Electrophysiological recordings on macaques demonstrate a segregation of the sensory responses within the claustrum (Remedios et al., 2010). The central part is responsive to auditory stimuli whereas the ventral part is responsive to visuals stimuli. The claustrum is thus described as multisensory but not as a multisensory integrator. Here, we also demonstrate visual activations in the ventral part of the claustrum in both audio-visual and purely visual contexts. Additionally, we report auditory activations in this region but not in the central claustrum as expected from Remedios and colleagues. Our observations of activation in the claustrum suggest that the perception of vocalisations is strongly modulated by the visual context and context-dependent visual association is based on semantic information in the absence of multisensory integration.

### Audio-visual association of social stimuli in the pulvinar

In contrast to the amygdala and claustrum, the contribution of the pulvinar to the processing of social auditory and visual stimuli is more complex. Specifically, we show a task-dependent sensory gradient such that auditory activations are located anteriorly, visual activations are located laterally and posteriorly and audio-visual activations are located medially. The medial pulvinar is the only sub-nucleus in which we could identify audio-visual integration, i.e. higher activations during bimodal, versus unimodal stimulation. Importantly, while anterior and medial pulvinar activations were driven by the task context set by the visual stimuli, this was not the case for the ventro-lateral pulvinar.

In Figure 8, we summarise the patterns of activation in the pulvinar. Several points are worth raising about these patterns. First, a global sensory gradient can be seen, with the anterior pulvinar being dedicated to auditory processing, the medial part to audio-visual and the lateral part of the nucleus to visual. Second, the dorso-medial lateral pulvinar (Pldm) is highly modulated by the task context and is only activated in a pure visual task and not in an audio-visual association task. Third, only the medial pulvinar and the ventral lateral pulvinar respond to visual social stimuli irrespective of context. It is also worth noting that this gradient does not follow the classical sub divisions defined based on its cytoarchitectonic properties, i.e. inferior pulvinar, lateral pulvinar and medial pulvinar (supplemental figures S3 and S4) (Walker, 1938; Olszewski et al., 1952; Gutierrez et al., 1995; Stepniewska and Kaas, 1997).

**Figure 8:**
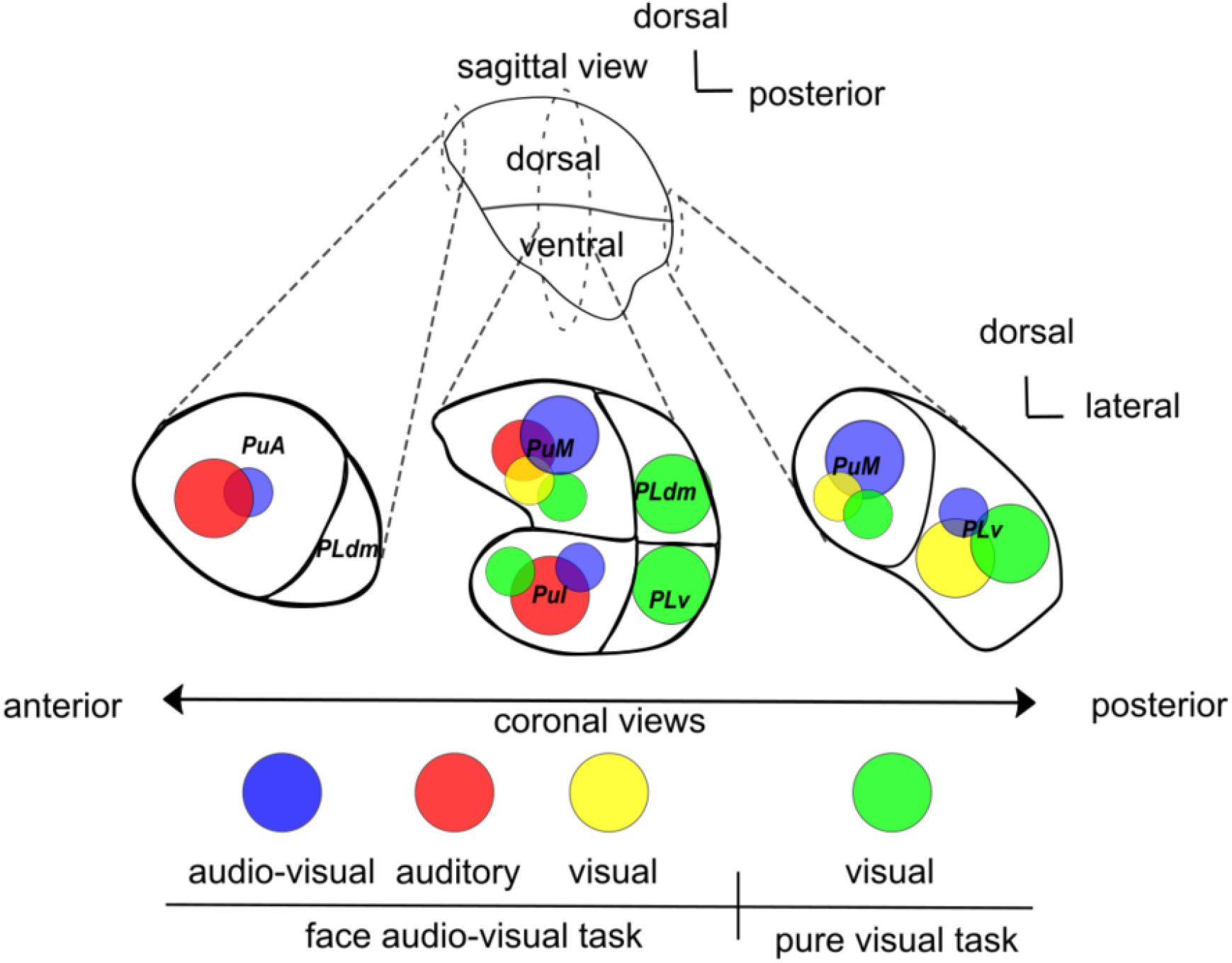
Summary of the activations within the pulvinar as a function of the task and sensory of stimulation. The circle size is largest in the regions of nucleus that present a higher activation. All circles correspond to significant activations or %SC.

### Pulvinar and Face perception

The lateral part of monkey pulvinar has been proved to contain face responsive neurons and the medial part is shown to be responsive to human facial expressions (Maior et al., 2010; Nguyen et al., 2013). Our observations from the pure visual task bring full support to this observation (figure 8, green circles). Using resting-state analysis, it has been found that the dorsal pulvinar, i.e. dorsal lateral and medial pulvinar, is functionally connected with face patches (Schwiedrzik et al., 2015). This is also the case for the ventral pulvinar. Generally speaking, the more anterior face patches connect to more anterior parts of the pulvinar thus defining an anterio-posterior functional connectivity gradient between the superior temporal sulcus and the pulvinar (Grimaldi et al., 2016). In addition, the stimulation of the two face patches AL and ML (respectively anterior lateral and medial lateral) of the superior temporal sulcus elicits activation in the inferior pulvinar (Moeller et al., 2008). Overall, this suggests that the entire pulvinar is potentially responsive to faces, these face responses being recruited differentially as a function of task and context.

Our study showed that a task implicitly calling for an association between auditory and visual social information, activates the medial as well as the ventro-lateral pulvinar. The pure visual task additionally activates the dorso-lateral and the inferior pulvinar. This is in agreement with the human pulvinar lesion literature, whereby a patient presenting with an entirely damaged unilateral pulvinar was not able to recognize fearful expressions in the contralesional field, while patients with damage limited to the anterior and lateral pulvinar showed no deficits in fear recognition (Ward et al., 2007). These results suggested fear recognition is mediated by the medial pulvinar. We propose to link these observations to the role of the pulvinar in emotional regulation, as discussed next. Additionally, the medial pulvinar is the unique pulvinar nucleus that presents audio-visual integration. This result supports the idea that this subregion implements multisensory integration (Froesel et al., 2021). As is the case for the amygdala, recent studies report multisensory integration in this brain structure. A macaque single cell recording study describes sub-additive and suppressive multisensory integration in the medial pulvinar (Vittek et al., 2022) while a recent human fMRI study of the pulvinar region describes multisensory enhancement during natural narrative speech perception (Ross et al., 2022). Here, we show that audio-visual integration can also be observed when the stimuli are matching only on semantic criterions and not classical multisensory matching. It follows the hypothesis formulated by Ross et al (2022) that multisensory enhancement in more naturalistic contexts involving understanding of semantics, recruits more than the typical multisensory network, notably the amygdala and the pulvinar. Given the fact that the medial pulvinar is highly connected with the limbic system and areas involved in the regulation of emotions such as anterior cingulate cortex, the temporal cortex, the temporo-parietal junction, the insula, the frontal parietal opercular cortex (Yeterian and Pandya, 1997; Rosenberg et al., 2009), together with the amygdala with which it is also connected (Jones and Burton, 1976), these structures are proposed to coordinate cortical networks to evaluate the biological significance of affective visual stimuli (Pessoa, 2010b). Their recruitment during audio-visual binding based on socioemotional context supports this hypothesis.

### Lateralisation

In the pulvinar, a processing bias in the left pulvinar has already been demonstrated in humans, which has been interpreted in the light of the role of the pulvinar in the attentional function (Padmala et al., 2010). In their monkey rsfMRI study, Schwiedrzik et al determined that the pulvinar, amygdala, hippocampus, caudate nucleus, claustrum and other sub cortical structures are functionally connected with face patches (Schwiedrzik et al., 2015). In addition, their left pulvinar activations were more widespread than the right activations. In our present study, activations during the pure visual context are exclusively identified in the left pulvinar. A left hemisphere bias for processing species-specific vocalisation was reported in a field study of rhesus monkeys (Hauser and Andersson, 1994). All this taken together with the fact that the SNR is not higher on the left that on the right (see supplemental figure S4), raises the possibility of functional lateralization bias in the monkey pulvinar. This will have to be further explored.

### Conclusion

In this study, we suggest that of the amygdala, claustrum, pulvinar play an essential role in multisensory integration of communicatively significant information. In addition, we demonstrate that there is a sensory audio-visual processing gradient within the pulvinar that is task dependent. We show that the medial pulvinar and amygdala, but not the claustrum, can be considered as audio-visual associators of social stimuli. We propose that these three sub cortical structures are part of an amodal network that modulates sensory perception as a function of the social context.

## Acknowledgements

S.B.H. were funded by the French National Research Agency (ANR)ANR-16-CE37-0009-01 grant and the LABEX CORTEX funding (ANR-11-LABX-0042) from the Université de Lyon, within the program Investissements d’Avenir (ANR-11-IDEX-0007) operated by the French National Research Agency (ANR). We thank Fidji Francioly and Laurence Boes for animal care, Franck Lamberton and Danièle Ibarrola for their MRI methodological support and Holly Rayson for their help on visual and auditory stimuli collection.

## Supplemental data

**Supplementary Table 1:**
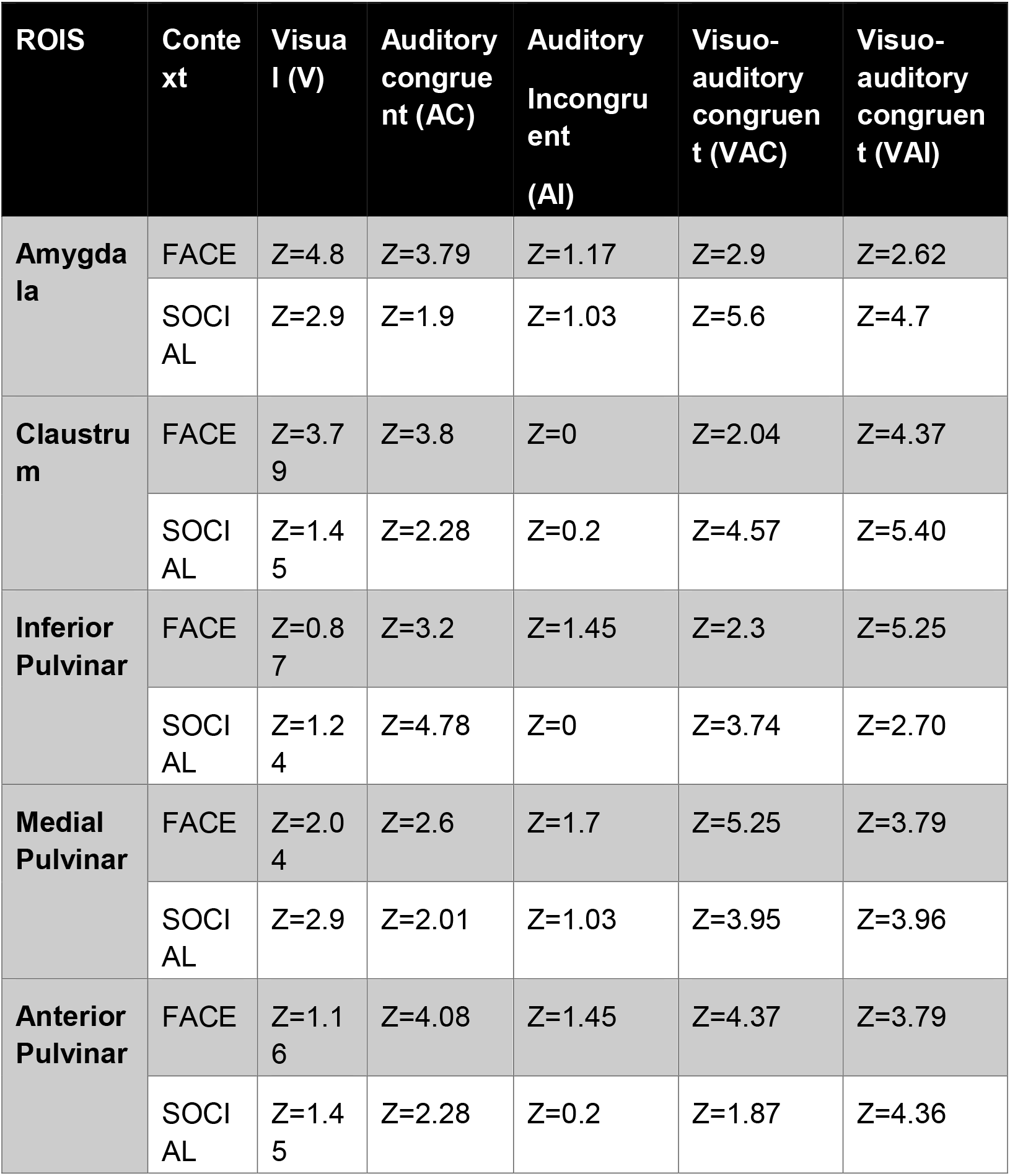
Effect sizes of the activations (percent signal change). Effect sizes of the Wilcoxon test performed on the percent signal changed. Significant results are presented in bold. This is a companion table to Figure 3.

**Supplemental figure S1:**
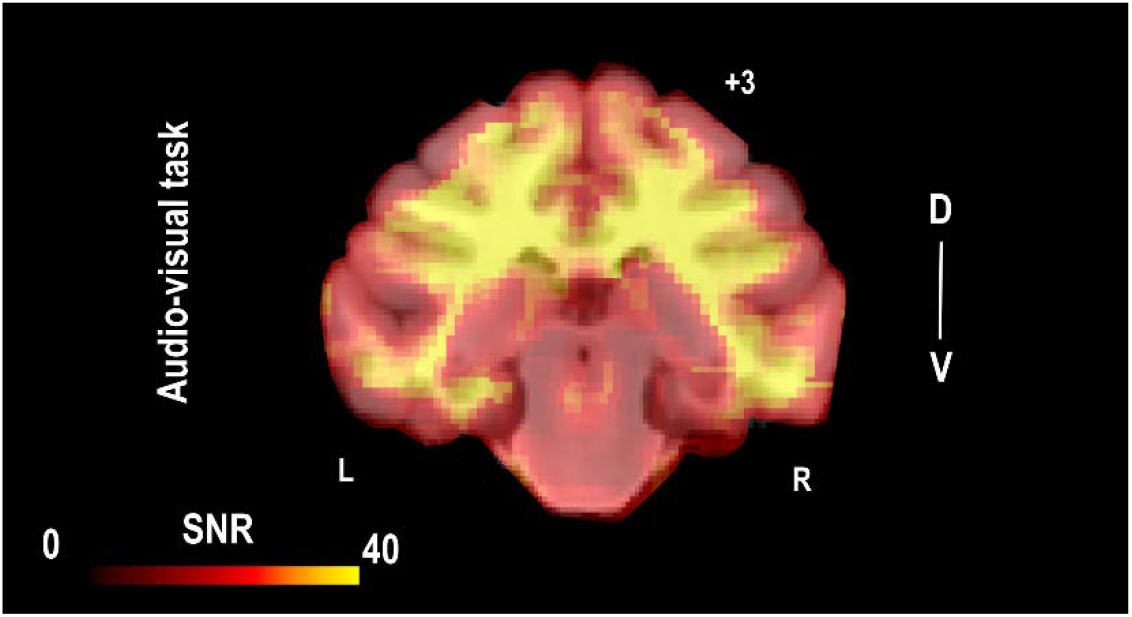
group SNR map for the audio-visual task.

**Supplemental figure S2:**
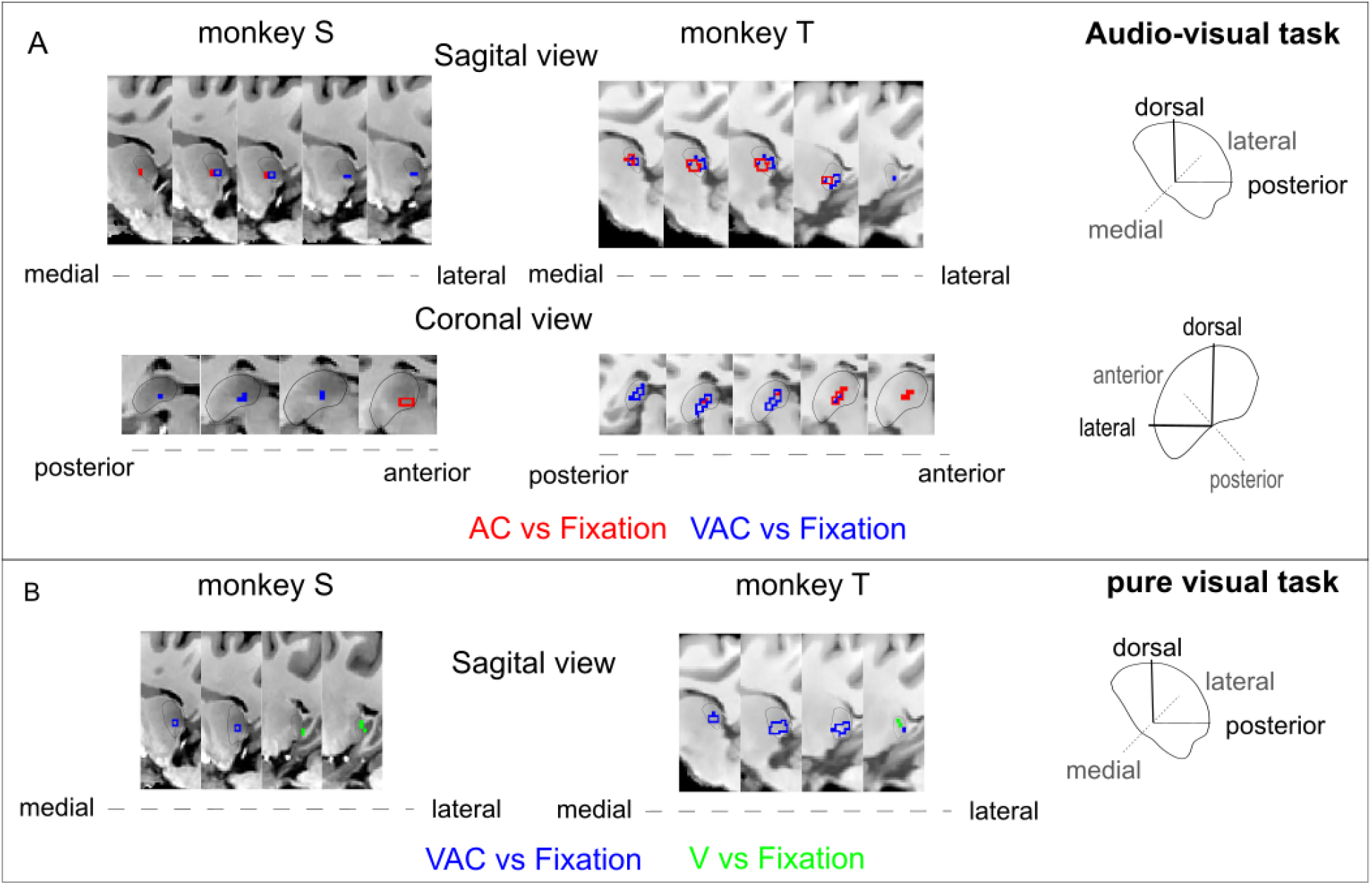
Pulvinar (and whole-brain) activations in FACE (F+ & F-) audio-visual and pure visual tasks per monkey. A) Whole-brain activation maps of the FACE context for both monkeys (monkey S and T), for auditory congruent (red) and audio-visual congruent (blue) contrasts vs. fixation. Axial, sagittal and coronal view are shown, zooming on the pulvinar. The activation outline in red and blue correspond to activation thresholds at the level of significance p<0.001 uncorrected, t-score 3.09, DF [1, 2600], and p<0.05 uncorrected, t-score 1.64 DF [1,2600] for monkey S. B) Whole-brain activation maps of the FACE context cumulated over both monkeys, audio-visual congruent condition of the audio-visual task (blue) and visual condition (aggressive + lipsmack faces) of the pure visual task (green). Note that even if the significance threshold for both monkeys are not the same, the gradient is present.

**Supplemental figure S3:**
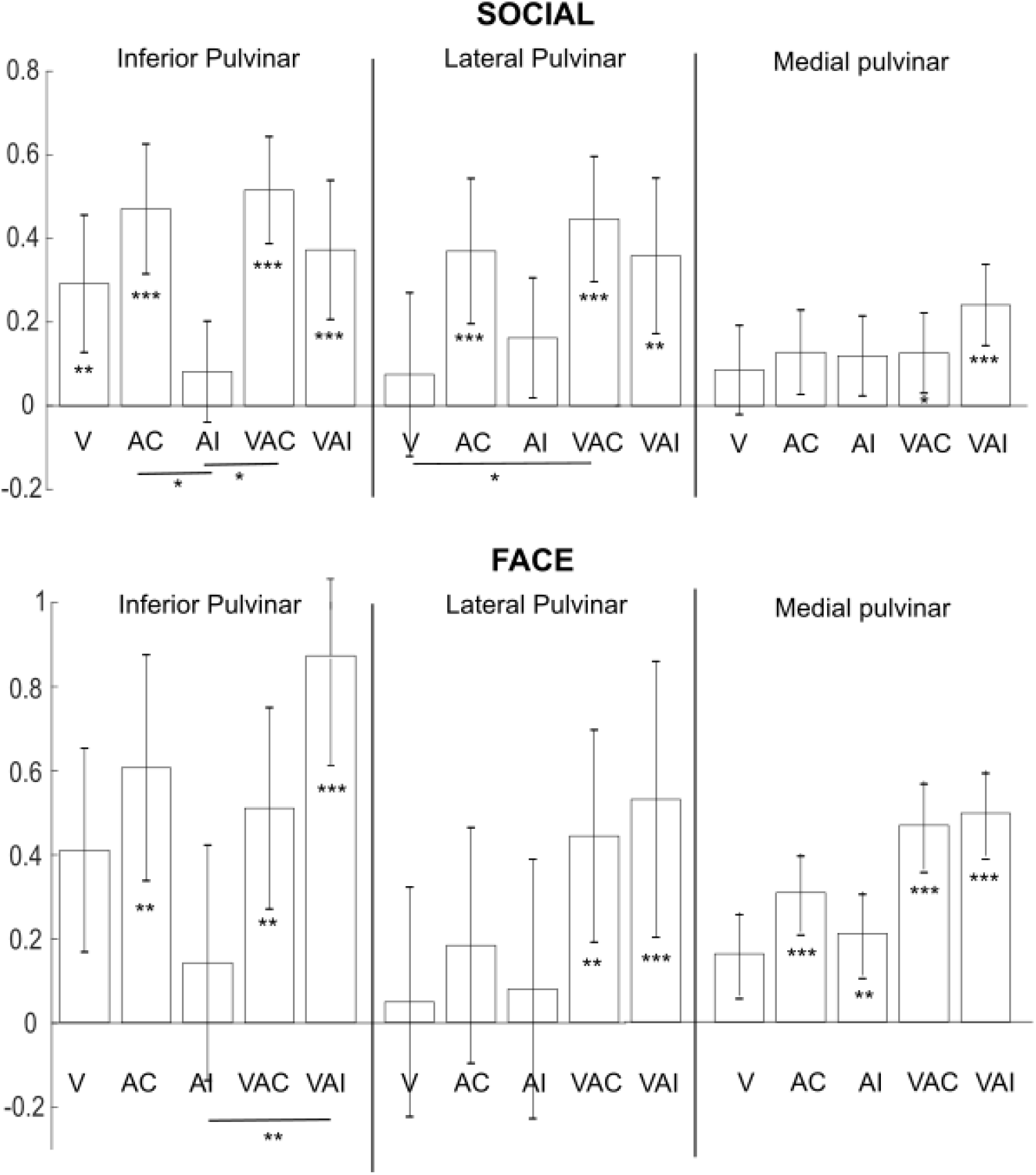
Percentage of signal change (%SC) in anatomical defined pulvinar ROIs for the audio-visual task for social and face contexts. Statistical differences relative to fixation and between conditions are indicates as follows: ***, p<0.001; **, p<0.01 (Wilcoxon nonparametric test). V: visual vs fixation, AC: auditory congruent vs fixation; AI: auditory incongruent vs fixation; VAC: audio-visual congruent vs fixation; VAI: audio-visual incongruent vs fixation. SOCIAL: PuI: V: Z=2.70, p=0.007; AC: Z=4.16, p<0.001; AI: 1.24, p=0.21, VAC: 5.4, p<0.001, VAI: 3.53 p<0.001. Friedman non-parametric test, X2_(4)_ = 16.34, p=0.0026, n=158, post hoc: VAC: Z=1.96, p=0.049, AC-AI: Z=2.48, p=0.013. PuL: V=0.8, p=0.4, AC: 3.32, p<0.001; AI: 1.87, p=0.06; VAC: 3.74, p<0.001; VAI: 2.9, p=0.0036. Friedman non-parametric test, X2(4) = 10.57, p=0.032, n=158, post hoc: V-VAC: Z=2.15, p=0.031. PuM: V: Z=1.39, p=0.29; AC: Z=1.45, p=0.14, AI: Z=1.66, p=0.09; VAC: Z=2.08, p=0.03; VAI: Z=3.53, p<0.001. Friedman non-parametric test, X2_(4)_ = 5.72, p=0.22, n=158. FACE: PuI: V: Z=0.87, p=0.38; AC: Z=3.2, p=0.0013; AI: 1.45, p=0.14; VAC: 3.21, p=0.0013, VAI: 5.25, p<0.001. Friedman nonparametric test, X2_(4)_ = 11.59, p=0.02, n=80, post hoc: V-AC: Z=1.96, p=0.049, AC-AI: Z=2.48, p=0.013. PuL: V=0.29, p=0.77; AC: 1.46, p=0.15; AI: 0.29, p=0.77; VAC: 3.21, p=0.0013; VAI: 3.5, p<0.001. Friedman non-parametric test, X2(4) = 8.6, p=0.07, n=80, post hoc: V-VAC: Z=2.15, p=0.031. PuM: V: Z=1.1, p=0.24; AC: Z=4.08, p<0.001; AI: Z=3.2, p=0.001; VAC: Z=4.9, p<0.001; VAI: Z=3.2, p=0.0013. Friedman non-parametric test, X2_(4)_ = 7.45, p=0.11, n=80.

**Supplemental figure S4:**
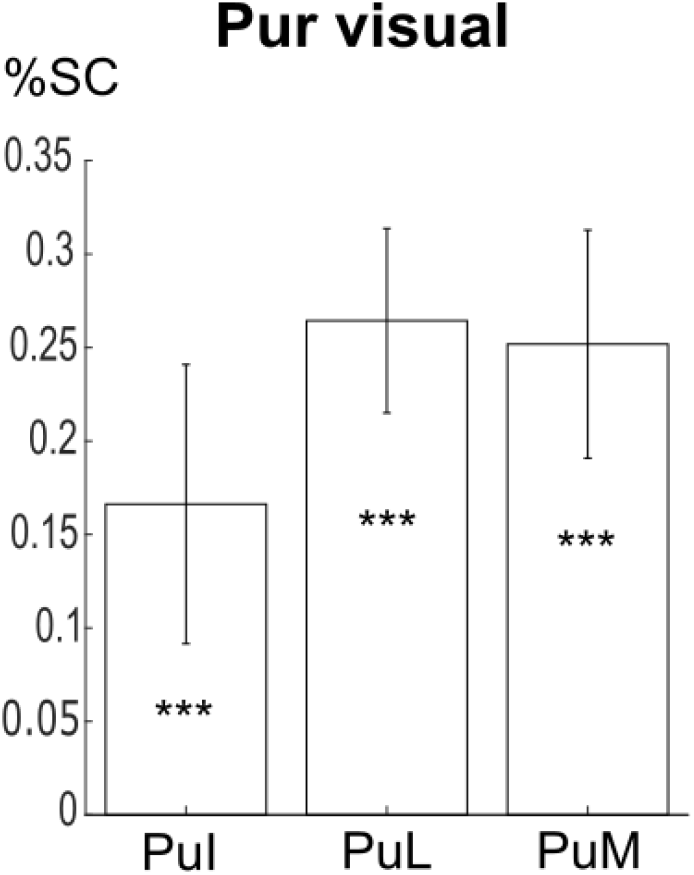
Percentage of signal change (%SC) in anatomical defined pulvinar ROIs for the pure visual task. Statistical differences relative to fixation and between conditions are indicates as follows: ***, p<0.001; **, p<0.01 (Wilcoxon non-parametric test: PuI: Z=3.99 p<0.001; PuL: 6.12, p<0.001; PuM: 3.73, p<0.001). PuI: inferior pulvinar; PuL: lateral pulvinar and PuM: medial pulvinar.

**Supplemental figure S5:**
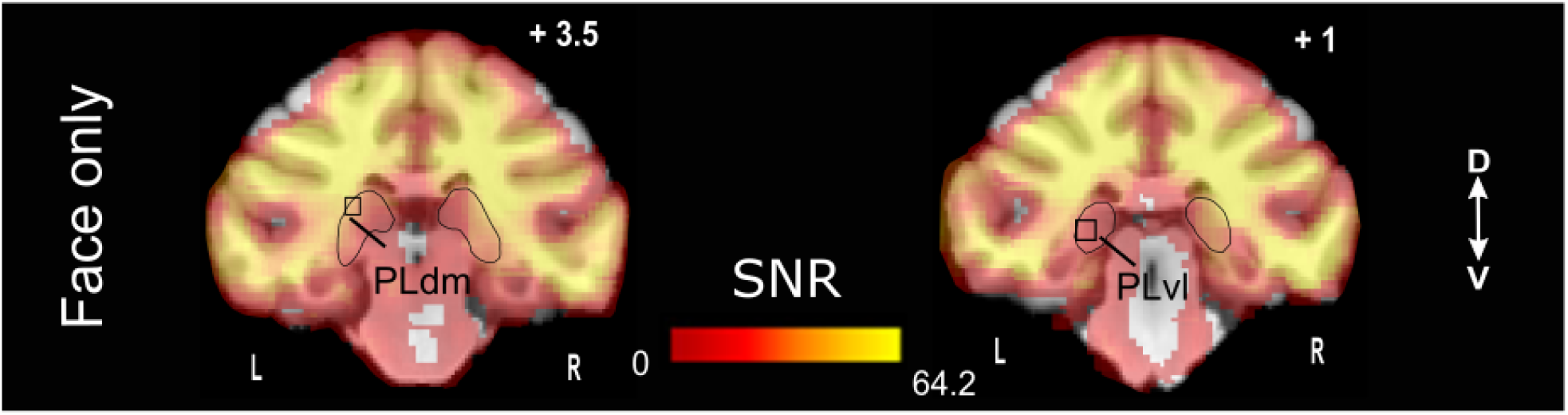
SNR map for the purely visual task both animals

**Supplementary Figure S6:**
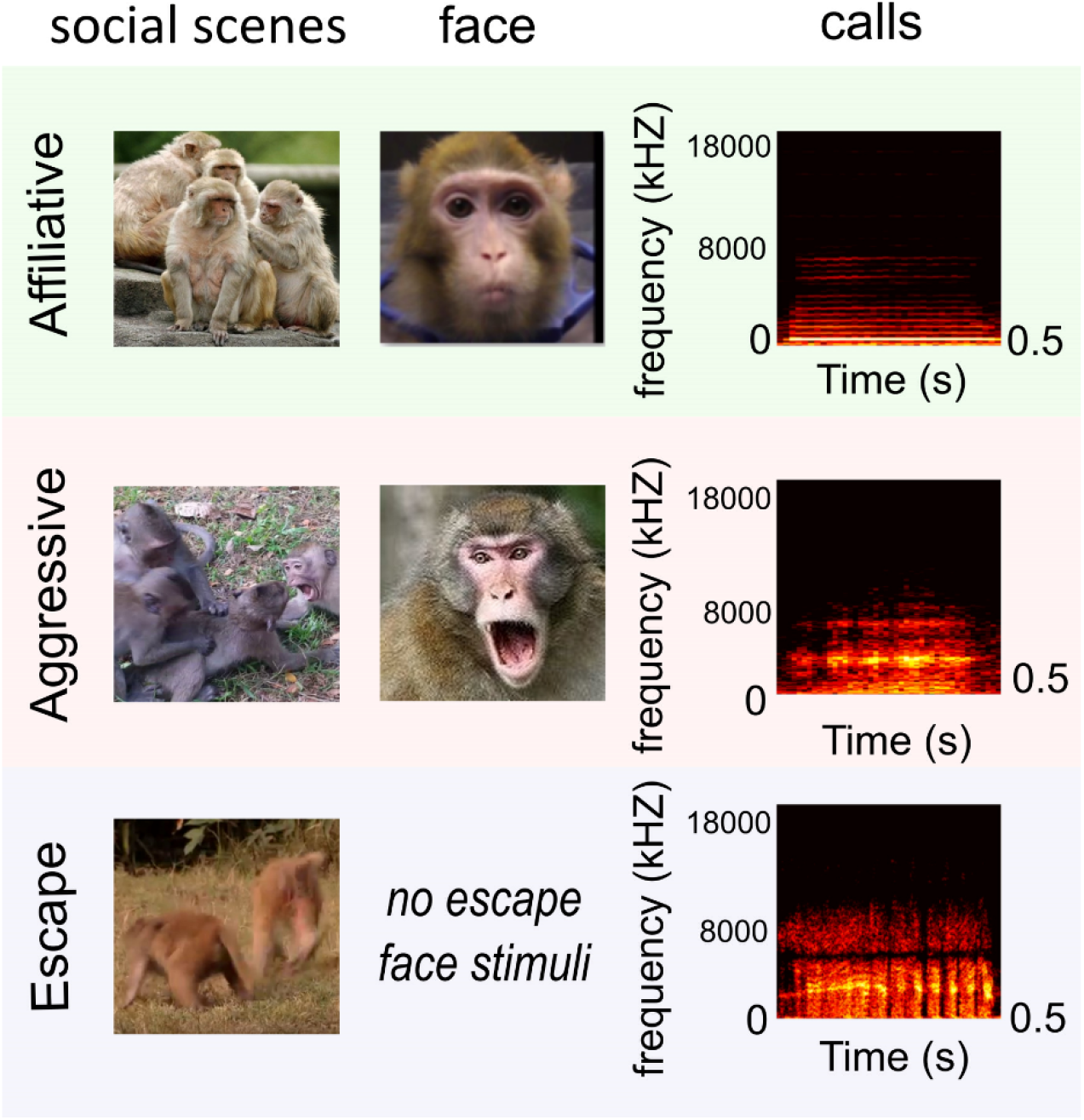
Example of visual and auditory stimuli. At the right are shown examples of visual stimuli used for social and face contexts. On the left, congruent calls spectrograms are associated to the visual stimuli are shown. The affiliative call is a coo, the aggressive congruent auditory stimulus is an aggressive call and the escape call is a scream. Stimuli were not strictly normalized in terms of in low visual and auditory feature properties, thus making their social meaning the dominant cue across the different stimuli of a given category. Stimuli were extracted from videos collected by the Ben Hamed lab, as well as by Marc Hauser on Cayo Santiago, Puerto Rico.

